# The novel compound AS1 abolishes second phase formalin nocifensive behavior

**DOI:** 10.1101/2025.02.06.636841

**Authors:** Arthur F. Brandao, Logan Condon, Kali Esancy, Benjamin Land, Ajay Dhaka

**Affiliations:** Department of Neurobiology and Biophysics, University of Washington, Seattle. United States of America; Department of Biochemistry, University of Washington School of Medicine, Seattle. United States of America; Department of Pharmacology, University of Washington School of Medicine, Seattle. United States of America; Center for the Neurobiology of Addiction, Pain, and Emotion, University of Washington School of Medicine, Seattle. United States of America; Graduate Program in Neuroscience, University of Washington, Seattle, United States of America

## Abstract

In larval zebrafish Analgesic Screen One (AS1), is a small molecule analgesic that reverses the valence of noxious stimuli rendering these aversive stimuli attractive. In this study, we evaluated the analgesic potential of AS1 in mice by measuring the effects of this compound on behavioral models of pain, locomotion and valence. We found that AS1 selectively abolished second phase nocifensive behavior elicited by the chemical irritant formalin, while having no effect on acute or inflammatory withdrawal thresholds evoked by thermal or mechanical stimuli. Unlike in larval zebrafish, AS1 did not induce attraction to aversive thermal stimuli. Neuronal activity profiling with the immediate-early gene c-Fos revealed that AS1 did not alter formalin evoked activity in the dorsal horn of spinal cord, but instead elevated activity in numerous brain regions associated with pain processing, analogous to findings in zebrafish. Further behavioral testing suggested that AS1 is not intrinsically aversive or appetitive but does promote reduced spontaneous locomotion and anxiogenic behavior. Collectively, these data suggest that AS1 may have select analgesic properties in mice.

## Introduction

Pain is the number one reason patients seek health care and greater than 20% of the US population is affected by chronic pain. Existing therapeutics have limited efficacy and a narrow therapeutic period, while also evoking deleterious side effects. Larval zebrafish have a number of attributes that lend themselves to inquiries into the biology of nociception. They are fully functioning animals, which must hunt for prey and assign appropriate valence to salient stimuli in order to survive. Importantly the spinal, subcortical and cortical structures that underpin pain sensation and the assignment of negative affect that drives the aversion and suffering evoked by painful stimuli are conserved in larval zebrafish (1–9). In an effort to identify novel analgesics, we utilized this species to perform a small molecule screen based on pain avoidance behavior. We identified analgesic screen 1 (AS1), a small molecule that reversed the valence of noxious stimuli, rendering them attractive to larval zebrafish in a dose dependent manner. AS1 was not appetitive on its own and mediated its effects independently of mu-opioid signaling. Furthermore, AS1 in the presence of noxious stimuli appeared to selectively engage neural correlates of the nucleus accumbens, striatum, hypothalamus, septal nuclei and the amygdala without affecting primary nociceptor activation. These areas are associated with affective pain processing and the assignment of valence to noxious stimuli in mammals. Intriguingly, the ability of AS1 to ascribe positive valence to noxious stimuli appeared to be separable from its analgesic properties (10).

Intrinsic to the emotional suffering elicited by pain perception is the attribution of a negative valence to nociceptive stimuli. A compound that could tune the valence of pain may prove to be a powerful therapeutic for the treatment of pain. To determine if the valence switching and/or analgesic properties of AS1 are conserved between larval zebrafish and mammals, we explored the effects of AS1 on behaviors elicited by noxious stimuli in mouse models of nociception. Here, we report that while the valence-switching properties of AS1 are not conserved, AS1 appears to act downstream of the spinal cord to selectively ablate second phase formalin-evoked nocifensive behaviors. Similar to zebrafish, AS1 increased neural activity, as measured by the expression of the immediate-early gene c-Fos, in brain areas that have been implicated in affective pain processing such as the shell of the nucleus accumbens, the anterior cingulate cortex, the paraventricular hypothalamus, and the paraventricular thalamus.

## Materials and Methods

### Animals

Female and male mice were separately group housed under controlled temperature at 20-26°C and 14-h light/10-h dark cycle. Polycarbonate cages (Allentown) with a cotton nesting square and with at least one enrichment material (igloo houses or paper tube) were used. Mice were given access to food and water ad libitum, and experiments were conducted in the light phase. All experiments were approved by the Institutional Animal Care and Use Committee (approved protocol #4216-01) University of Washington. All experiments were performed with the investigator blinded to treatment.

### AS1 preparation

The AS1 (20mg/ml) was first dissolved in 100% dimethyl sulfoxide (DMSO, 1mL. Fischer Scientific. Fair Lawn, NJ), aliquoted in 100µl and stored at -20°C. On experimental days, aliquots of AS1 were defrosted and suspended in DMSO and Cremophor-EL® (#238470, Millipore. Temecula, CA), and diluted with 0.9% (NaCl) saline to the final working solution (2.5mg/kg, 3.75mg/kg, or 5mg/kg). The vehicle solution was prepared at the same time and concentration of the AS1 solution (5% DMSO, 5% Cremophor, and 90% saline). AS1 or vehicle was intraperitoneal injected (i.p.) at 10µl/g final dose.

### Formalin assay

A 2.5% formalin solution was made by diluting 25µl of the stock formalin (36.5-38%; #F8775, Sigma, Aldrich) in 975µl of saline (NaCl 0.9%). Then, 20µl of formalin solution was injected into the plantar surface of the left hind paw of the mice by using an insulin (31-gauge needle - Medline) or a Hamilton syringe (25µl, #702 LT SYR) with a 30-gauge needle connected. For the formalin test, mice were habituated for 60 minutes in a Plexiglass box (9cm x 11.5cm x 14cm) atop an acrylic floor in a quiet and temperature-controlled room. Twenty minutes prior to the formalin injection into the left hind paw, each mouse was separately and carefully removed from the apparatus for the AS1 or vehicle (i.p.) injection. Mice were then subsequently returned to the apparatus. After 20 minutes, each mouse was again separately and carefully removed from the apparatus and given an intraplantar injection of formalin. All sessions were video recorded for 45 minutes. The formalin-induced licking behavior time was recorded in 5-min bins.

### Hotplate Assay

The hotplate assay was performed essentially as previously described (11). Mice were first habituated to the testing room for at least 60 minutes. The hot plate test was conducted in a solid heat plate (27.5 cm x 26.3 cm x 1.5 cm, IITC Life Science, Los Angeles, CA) with a Plexiglas box on top, and temperature was set to 57°C ±0.5 °C. The trial was video recorded (Canon HD, Vixia HF200) and the thermal nocifensive behaviors were measured by the latency time to lick the paws or jump. Behavior was measured at baseline (no treatment) and 20, 50 and 110 minutes after treatment. The AS1 (5mg/kg) or vehicle were administrated 10 minutes after the baseline time point. A 30 second cutoff time was used to avoid tissue damage. To assess thermal hyperalgesia, Complete Freund’s adjuvant (CFA, Sigma) was injected in the left hind paw (10µg in 20µl diluted 1:1 in saline 0.9% NaCl). Mice were assessed as described above 20 minutes after AS1 (5mg/kg) or vehicle treatment at 0, 1, 3 and 7 days after CFA administration.

### Tail-flick Assay

The tail-flick test was performed in a hot water bath with defined temperature at 52.5°C ± 0.5 °C (Thermo Scientific Precision #2833). Mice were habituated to the testing room for at least 60 minutes prior to the test. Each mouse was carefully removed from their cage and gently restricted using a soft cloth. Then, 2.5cm of the tip tail was submersed in the hot water until a nociceptive response of tail withdrawal was observed. The tail-flick latency was manually registered using a stopwatch and maximum tail reaction time of 15 seconds was used as a cut-off to avoid tissue damage. The nociceptive tail response was measured at baseline (no treatment) and 20, 50 and 110 minutes after treatment. The AS1 (5mg/kg) or vehicle were administrated 10 minutes after the baseline time point.

### Thermal Place Preference

The thermal place preference assay was conducted in a dual solid-state control/heated plate (34x33cm, AHP-1200°CP, Teca) where each mouse freely explored both sides of the plate. A white acrylic rectangular arena (35cm width, 34cm length, and 15cm height) was placed on top of the control/heated plate, which created four individual rows of control/heat place preference. The control side temperature of the plate was fixed at 30°C ±0.5 °C and the heated side was fixed at 45°C ±0.5 °C. A second temperature probe was used to fine adjust plate temperature. Mice were first habituated in the indirect-light experimental room in their cages for 60 minutes. The mice were then injected with 5mg/kg AS1 dose or vehicle (i.p.) 20 minutes prior to being placed at the center point of each row on the control/heated plate. Each trial consisted of mice freely exploring both sides of the plate for 20 minutes. Each trial was video recorded (Canon HD, Vixia HF200) and animal position was determined utilizing tracking software (EthoVision XT, Nodus Technology Inc.).

### Cold Plantar Assay

A cold plantar test probe was generated using a cut off 5ml syringe filled with freshly crushed dry ice (12). Prior to the assay, mice were acclimated on Plexiglass box (9cm x 11.5cm x 14cm) atop a glass floor of 4.75mm for 60 minutes. The cold nociception response was assayed at baseline (no treatment) and 0, 20, 50, 80, and 110 minutes after treatment. The AS1 (5mg/kg) or vehicle were administrated (i.p.) 60 minutes after the baseline time point. The test probe was applied to the glass floor directly underneath a hind paw. Both right and left hind paws withdrawal latencies were recorded and averaged. At least 3min were allowed between consecutive trials.

### Inflammatory Allodynia

Mice were first placed for 60 minutes in Plexiglass box (9 cm x 11.5 cm x 14 cm) with wire grid floors in a quiet and temperature-controlled room for habituation. Complete Freund’s adjuvant (CFA, Sigma) was injected in the left hind paw (10µg in 20µl diluted 1:1 in saline 0.9% NaCl) and the left hind paw was measured. Paw withdrawal threshold (PWT) was assessed pre- and post-treatment of AS1 (5mg/kg) or vehicle with a calibrated series of von Frey filaments. The PWT measurements were performed at baseline (without CFA) and 1, 3, and 7 days after CFA injection. The AS1 or vehicle was injected 20 minutes prior to PWT measurement. The ascendent stimulus method was used to estimate the PWT (13–15).

### Conditioned Place Preference

The conditioned place preference test (CPP) was adapted from (11). Briefly, the CPP test was conducted in a sound-protected chamber with a Plexiglass box inside divided with two distinct textural sides (metal mesh flooring versus a parallel metal bar flooring cues). The chamber was equipped with an overhead camera to track mouse position and mice-side preference was determined via EthoVision XT software (Nodus Technology Inc.). Mice were habituated to the testing room for at least 60 minutes before each day of test and the test was performed on 5 consecutive days. In the pre-test (day 1), mice were allowed to freely explore both sides of the box for 15 minutes and the total time spent on each side of the box was calculated. Only mice that did not present a strong baseline preference (<60% of the total time) in the pre-test for any side were used for the analysis of the CPP test (>85% mice met the criteria). In the next 3 consecutive days (day 2, 3, and 4), mice were conditioned in 2 pairing sessions each day. In the morning sessions, mice were given saline solution (i.p.) and paired with one of the sides. In the afternoon sessions, mice were given the AS1 (5mg/kg, i.p.) or vehicle and paired with the opposite side. The AS1 or vehicle were injected 20 minutes before mice were placed in the chambers and mice were conditioned for 60 minutes. In the test day (day 5), mice were allowed to freely explore both sides of the box exactly as on day 1 and the total time spent on each side was analyzed. The preference score was analyzed and calculated by the time expended on the drug-paired (AS1 or vehicle) side minus the time expended in the control side.

### Open Field

The open field test (OFT) was performed in a white square plastic box (40x40x30cm) in an 80-90 lux lighting and overhead camera to track mouse movement was used. For the OFT, mice were first habituated to the testing room for at least 60 minutes and injected with AS1 (5mg/kg) or vehicle 20 minutes prior to the test. Mice were placed at the center of the OF box and monitored for 30 minutes. The total time spent in the center (20x20cm, 50% of total box area) and in the border of the OF box, the total distance traveled, and mean velocity were analyzed with EthoVision XT software (Nodus Technology Inc.).

### Intrathecal Injection

One day prior to the intrathecal (i.t.) injection, mouse back fur was shaved to facilitate localization of the L4-L6 vertebrae. For i.t. injections, a 25µl Hamilton syringe connect to a 30-gauge needle was used to insert and deliver the solution into the L5 or L6 spinous process interverbal space (16). AS1 (5µg/µl, 5µl) or vehicle (5µl) were injected 20 minutes prior to the intraplantar formalin (2.5%) injection. Formalin-induced licking behavior was assessed as described above.

### Immunohistochemisty

Mice were transcardially perfused with 1X phosphate buffered saline (PBS) and 4% paraformaldehyde (PFA). Brains or spinal cord were extracted, post-fixed in PFA solution at 4 °C overnight, and cryoprotected in 30% sucrose solution in PBS. Isopentane (Sigma Aldrich) and embedding medium for frozen tissue (Tissue-Tek, Sakura) were used to freeze all brains. Tissue was cut into 40µm (brain) or 30µm (spinal cord) coronal sections using a cryostat (Leica CM1850). For the cFos immunostaining, the sections were blocked with PBS containing 5% normal donkey serum (NDS) and 0.3% Triton X-100 for 1 h at room temperature. Sections were incubated in blocking buffer (5% NDS in 0.3% Triton X-100) at 4 °C for 24 hours with cFos primary antibody (dilution 1:500, rat monoclonal, Synaptic System #226-017, Goettingen, Germany). After three washes (10 min each) in PBS, sections were incubated with Alexa-conjugated secondary antibodies against the corresponding species (1:500, Alexa-Flour 647 anti-rat, # 712-606-150 Jackson ImmunoResearch Laboratories Inc., West Grove, USA) for two hours at room temperature. Sections were once again washed in PBS three times and were mounted and covered using Antifade Vectashield Mounting Medium with DAPI (Vector Laboratories. Burlingame, CA). Mounted slides were imaged on a fluorescent Keyence microscope (BZ-X800). Stained brain sections were quantified using an open source bioimage analysis software (QuPath) and Fiji/ImageJ (17–19). Three sections per brain structure and both right and left sides of brain hemisphere were quantified by researchers blinded to the experimental conditions. We analyzed the anterior cingulate area (ACA), nucleus accumbens core (NAc-Core) and shell (NAc-Shell), basolateral (BLA) and central (CeA) amygdala, paraventricular nucleus of thalamus (PVT), paraventricular hypothalamic nucleus (PVH), lateral septal nucleus (LS), and primary motor area (MOp). Stained spinal cord sections were manually quantified by researchers blinded to the experimental conditions.

### Data Analysis

Data are presented as mean and standard error of the mean (SEM). Statistical tests were performed using GraphPad Prism. The Shapiro-Wilk test was conducted, and a parametric test was applied when data normality distribution was reached. The RM-ANOVA were followed by post-hoc comparison using Bonferroni test. The Mann-Whitney U-test with continuity correction was applied to assess statistical significance of groups in non-parametric data. Paired and unpaired Student’s t test was used to compare two groups when necessary. Differences were considered significant at p<0.05 (* p<0.05; **p≤0.01; ***p≤0.001).

## Results

### AS1 abolishes second phase formalin nocifensive behavior

When injected into the hind paw of rodents, formalin evokes a biphasic pain response, characterized by nocifensive behaviors such as the biting, licking, lifting and flicking of the affected paw. This test is often used to evaluate the analgesic potential of compounds as it allows for the assessment of spontaneous nocifensive behavior in freely moving animals (20–22). We performed dose-dependent analysis to investigate whether AS1 (2.5, 3.75, and 5 mg/kg) treatment altered formalin-induced nocifensive behavior. In all cases, AS1 was administered intraperitoneally 20 minutes prior to formalin. None of these doses appeared to cause gross deficits in motor function. At the 2.5 mg/kg dose, there was no effect on formalin-evoked responses at any time interval as reflected in cumulative time engaged in nocifensive behavior in either the first (AS1: 114.60 ±9.48 vs Vehicle: 131.90 ±19.58 sec, p=0.35) and second (AS1: 492.70 ±42.23 vs Vehicle: 397.75 ±57.09 sec, p=0.28) phases (Fig 1A and 1B). At the 3.75 mg/kg dose, we observed a delay in second phase formalin-induced licking behavior, with a reduction of nocifensive behavior at the 20min (AS1: 49.71 ±10.47 vs Vehicle: 141.75 ±12.67 sec, p<0.001) timepoint but an increase in nocifensive behavior time at the 40 min timepoint when compared to vehicle treated animals (AS1: 42.42 ±9.68 vs Vehicle: 6.33 ±3.76 sec, p<0.05) timepoint (Figure 1D). However, again there was no statistical difference in cumulative first (AS1: 103.78 ±9.78 vs Vehicle: 132.16 ±13.33 sec, p=0.12) or second (AS1: 390.85 ±43.19 vs Vehicle: 498.83 ±31.46 sec, p=0.095) phase behavior (Fig1C). Strikingly at the 5 mg/kg dose, we observed a near total abolition of second phase nocifensive behavior (AS1: 34.14 ±16.65 vs Vehicle: 550.26 ±59.36 sec, p<0.001) with only a moderate effect on first phase behavior (AS1: 75.47 ±7.46 vs Vehicle: 122.51 ±15.61 sec, p<0.05) (Fig 1F). Using pos-hoc testing, we observed several timepoints where AS1 significantly reduced formalin-evoked nocifensive behavior: 20min (AS1: 3.48 ±2.56 vs Vehicle: 133.88 ±17.76 sec, p<0.001), 25min (AS1: 1.85 ±1.06 vs Vehicle: 141.24 ±17.41 sec, p<0.001), and 30min (AS1: 2.89 ±1.58 vs Vehicle: 145.48 ±23.74 sec, p<0.01) timepoints, (Fig 1E). These data suggest that AS1 may suppress formalin-evoked behavior with a tight dose-response curve and that the effects of AS1 in this assay persist until at least 60 minutes post-administration. Since only the 5 mg/kg dose of AS1 had a dramatic effect on formalin induced nociceptive behavior, we employed this dosage for subsequent investigations into its anti-nociceptive properties.

**Fig 1.**
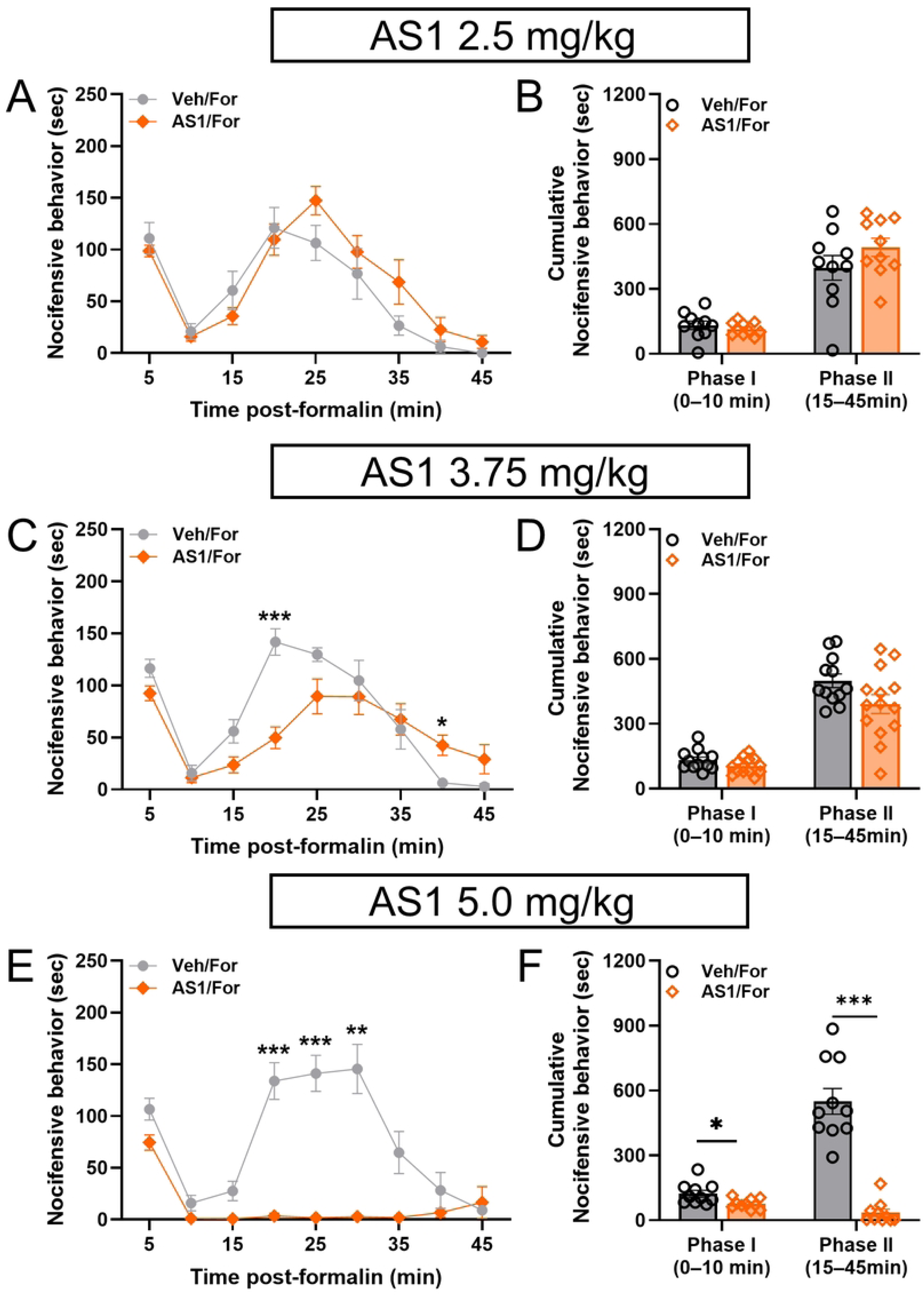
AS1 inhibits second phase formalin-induced nocifensive behavior. (A,C,E) Time course of formalin evoked nocifensive behavior. (C,D,F) Total response time for phase one and phase two formalin-evoked nocifensive behavior (A,B) AS1 (2.5mg/kg) has no effect on formalin-evoked nocifensive behavior (Veh/For (n=10/5 females), AS1/For (n=10/5 females). (C,D) AS1 (3.75mg/kg) alters specific timepoints (20 min, 40 min) but has no effect on cumulative formalin evoked nocifensive behavior (Veh/For (n=12/6 females), AS1/For (n=14/6 females). (E,F) AS1 (5.0mg/kg) abolishes second phase formalin responses (Veh/For, n=10/5 females), AS1/For (n=10/5 females).*p<0.05, **P<0.01, ***p<0.001. Veh=Vehicle, For=formalin. A,C, E, RM-ANOVA with Bonferroni’s multiple comparison. B,D, F, Unpaired student’s t-test.

### AS1 has no effect on withdrawal responses to thermal and mechanical stimuli in the presence or absence of inflammation

We next investigated whether AS1 (5mg/kg) would affect other acute nociceptive/withdrawal responses. We assessed the effects of AS1 on acute thermal nociceptive responses using the hot plate (57°C) and tail flick (52°C) tests (11,15). For the hotplate assay, we measured the latency to first nocifensive response, before treatment and 20-, 50- and 110-minutes post-treatment. Pretreatment baselines were subtracted from each timepoint to account for any group variance. We found no differences between AS1 and vehicle treated groups. The RM-ANOVA did not reveal an effect of treatment (F(1, 22)=0.06, p=0.81) (**Fig 2A**).

**Fig 2.**
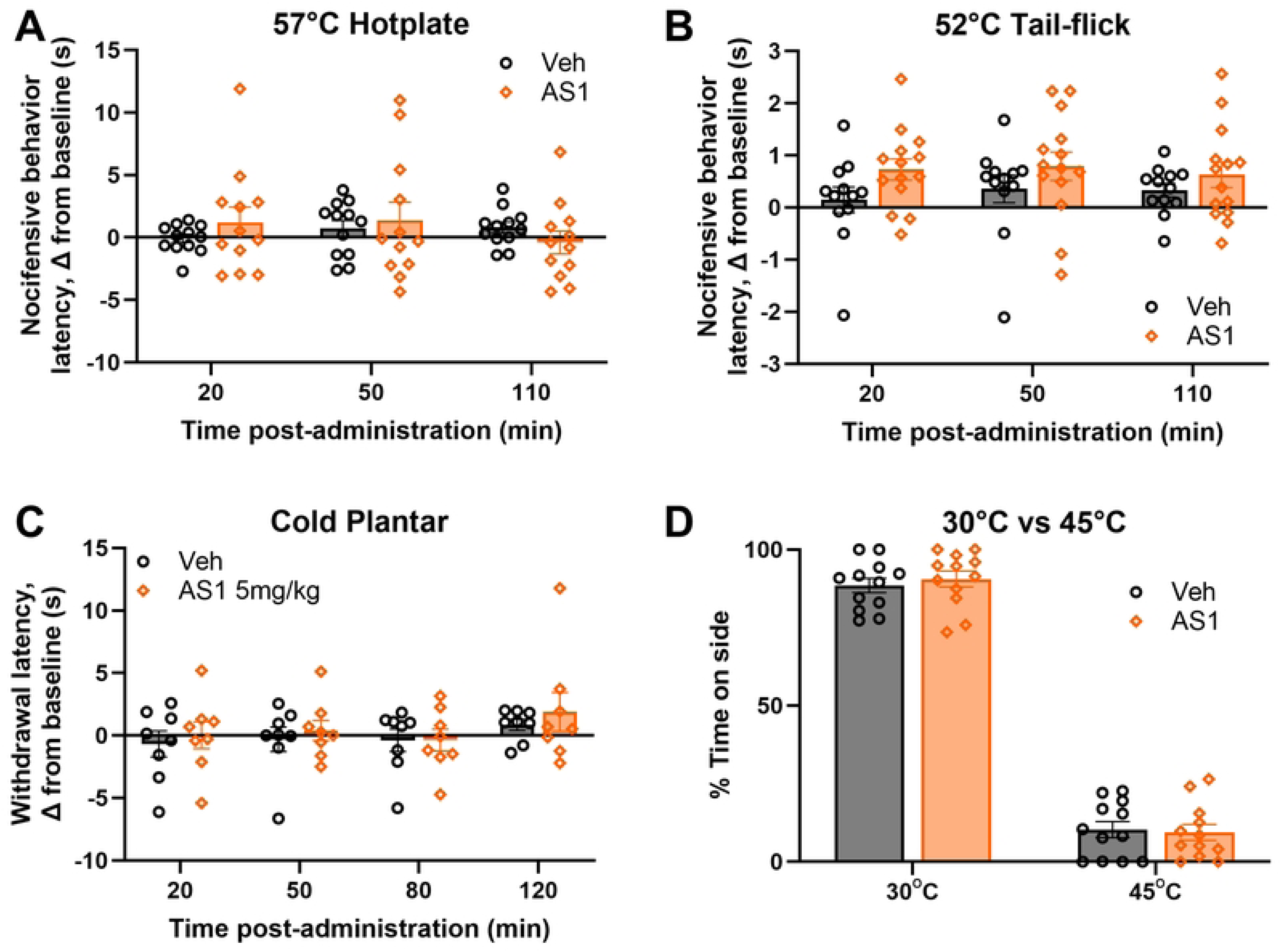
AS1 does not alter acute withdrawal responses or aversion to temperature. (A) AS1 (5mg/kg) did not alter nociceptive response latency in Hotplate (57^ο^C) test (Veh n=12/7 females, AS1 n=12/6 females). (B) AS1 (5mg/kg) had no effect on tail-flick response latency (Veh n=12/6 females, AS1 n=14/7 females). (C) AS1 (5mg/kg) had no effect on cold plantar withdrawal latency (Veh n=8/4 females, AS1 n=8/4 females). (D) AS1 (5mg/kg) did not affect thermal place preference (30^ο^C vs 45^ο^C) (Veh n=12/6 females, AS1 n=12/7 females). Veh=Vehicle, For=formalin. A,B,C, RM-ANOVA with Bonferroni’s multiple comparison. D, Unpaired student’s t-test.

We next measured heat withdrawal latency with the tail flick assay which unlike the hot plate, is not thought to require supraspinal circuitry (15). Again, pretreatment baselines were subtracted from each timepoint to account for any group variance. We found no differences between AS1 and vehicle treated groups when assessing tail flick responses. The RM-ANOVA test did not reveal a main effect of treatment (F(1, 24)=3.68, p=0.12) (**Error! Reference source not found.**).

To assess withdrawal responses to cold temperature we employed the cold plantar assay (12). AS1 (5mg/kg) had no effect on the latency response to a cold plantar stimulus. As above, pretreatment baselines were subtracted from each timepoint to account for any group variance. Post hoc analysis revealed no differences from vehicle treatment at any timepoint and the RM-ANOVA test did not reveal a main effect of treatment (F(1, 56)=0.74, p=0.39) (Fig 2C).

To determine if AS1 altered mechanical sensitivity, we measured hind paw withdrawal thresholds using von Frey filaments. We assessed withdrawal thresholds after vehicle or AS1 (5mg/kg) administration under baseline conditions and 1-, 3- and 7-days after CFA induced inflammation. Pretreatment baselines from prior to AS1 or vehicle treatment taken on the baseline day were subtracted from each timepoint to account for any group variance. RM-ANOVA revealed no difference in pre-CFA baseline paw withdrawal thresholds between AS1 and vehicle treated animals. Furthermore, while analysis revealed inflammatory mechanical allodynia 1- and 3-days post CFA administration in both AS1 and vehicle treated groups, there was no difference between AS1 and vehicle treated animals at any timepoint following CFA (Fig 3A). Taken together these data indicate that at this dosage AS1 had no effect on mechanical withdrawal thresholds acutely or after inflammation.

**Fig 3.**
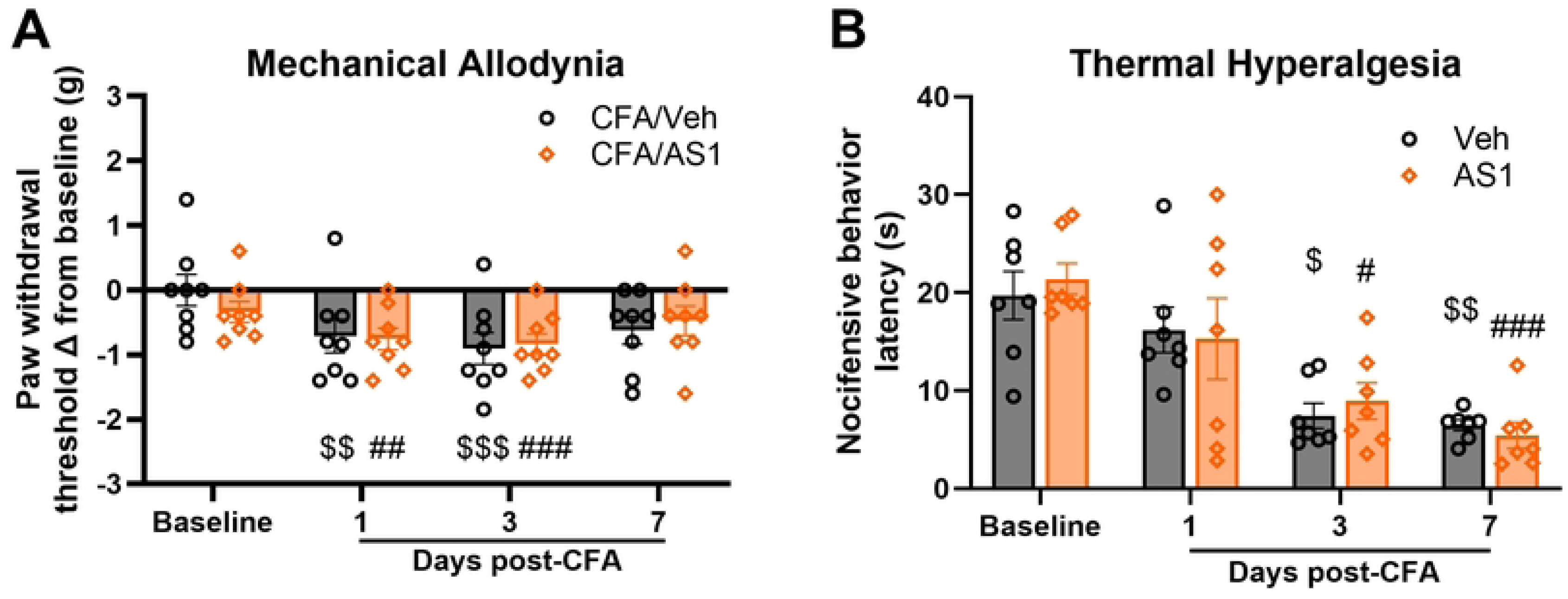
AS1 had no effect on mechanical allodynia and thermal hyperalgesia induced by CFA. (A) CFA induced mechanical allodynia 1- and 3-days post CFA. PWTs of AS1 (5mg/kg) treated animals did not differ from vehicle treated animals at any timepoint (CFA/Veh, n=8/4 females, CFA/AS1, n=8/4 females). (B) CFA induced thermal hyperalgesia 3- and 7-days post CFA. Nocifensive behavior latency did not differ between AS1 (5mg/kg)- and vehicle-treated animals at any timepoint (CFA/Veh, n=7/3 females, CFA/AS1, n=7/3 females). ^$,#^p<0.05, ^$$,##^P<0.01, ^$$$,###^p<0.001. A,B, Veh=Vehicle, For=formalin. RM-ANOVA with Bonferroni’s multiple comparison. $ denotes comparison with vehicle-treated baseline. # denotes comparison with AS1-treated baseline. A,B RM-ANOVA with Bonferroni’s multiple comparison.

We also examined if AS1 could alter thermal hyperalgesia evoked by CFA utilizing the hotplate test. Again, we found that AS1 did not affect the latency of withdrawal responses at baseline or after CFA-evoked hyperalgesia 1-,3- and 7-days after induction (Fig 3B). Taken together these data suggest that AS1 (5mg/kg), a dose that abolishes second phase formalin responses, has no effect on acute and inflammatory withdrawal responses to thermal and mechanical stimuli.

### AS1 does not alter noxious heat aversion

Since AS1 was found to reverse thermal place aversion in larval zebrafish, rendering aversive temperatures attractive, we explored whether AS1 might have similar properties in mice (10). We tested mice in a 30°C versus 45°C temperature choice assay to assess temperature preference after vehicle or AS1 (5mg/kg) administration. These temperatures were chosen since 45°C drives robust aversion in mice. Unlike in zebrafish AS1 had no effect on thermal aversion. As measured by percent time spent in each zone, there was no difference between AS1 and vehicle treated animals with both groups robustly avoiding the 45°C zone (AS1: 30°C zone, 90.6±2.5 % time; 45°C zone, 9.4±2.5 % time vs Vehicle: 30°C zone, 88.5±2.3 % time; 45°C zone, 10.3±2.6 % time, p=0.5 (30°C), p=0.8 (45°C)) (Fig 2D). This data suggests the ability of AS1 to induce attraction to aversive stimuli is not conserved in mice nor does it alter heat aversion.

### AS1 does not appear to be appetitive or aversive

In larval zebrafish, AS1 did not appear to be appetitive or aversive nor did it appear to engage mu opioid signaling (10). Furthermore, analgesic compounds, notably mu opioids, can have rewarding properties (23). To determine if AS1 (5mg/kg) is appetitive or aversive on its own, we performed a conditioned place preference assay. A paired T-test revealed no difference in the percentage of time expended on drug-paired side for either vehicle (pre 41.93 ±2.38 vs 41.31 ±2.77%, p=0.72) or AS1 (pre 42.88 ±1.53 vs 44.10 ±2.52%, p=0.60) treated mice when comparing the pre-test to the post-test (Fig 4A). Furthermore, there was no difference in the preference score of vehicle or AS1 treated animals (Veh -5.11 ±15.42 vs 11.47 ±20.82 sec, p=0.53) (Fig 4B). Taken together these data indicates that this dose of AS1 does not induce place preference or aversion.

**Fig 4.**
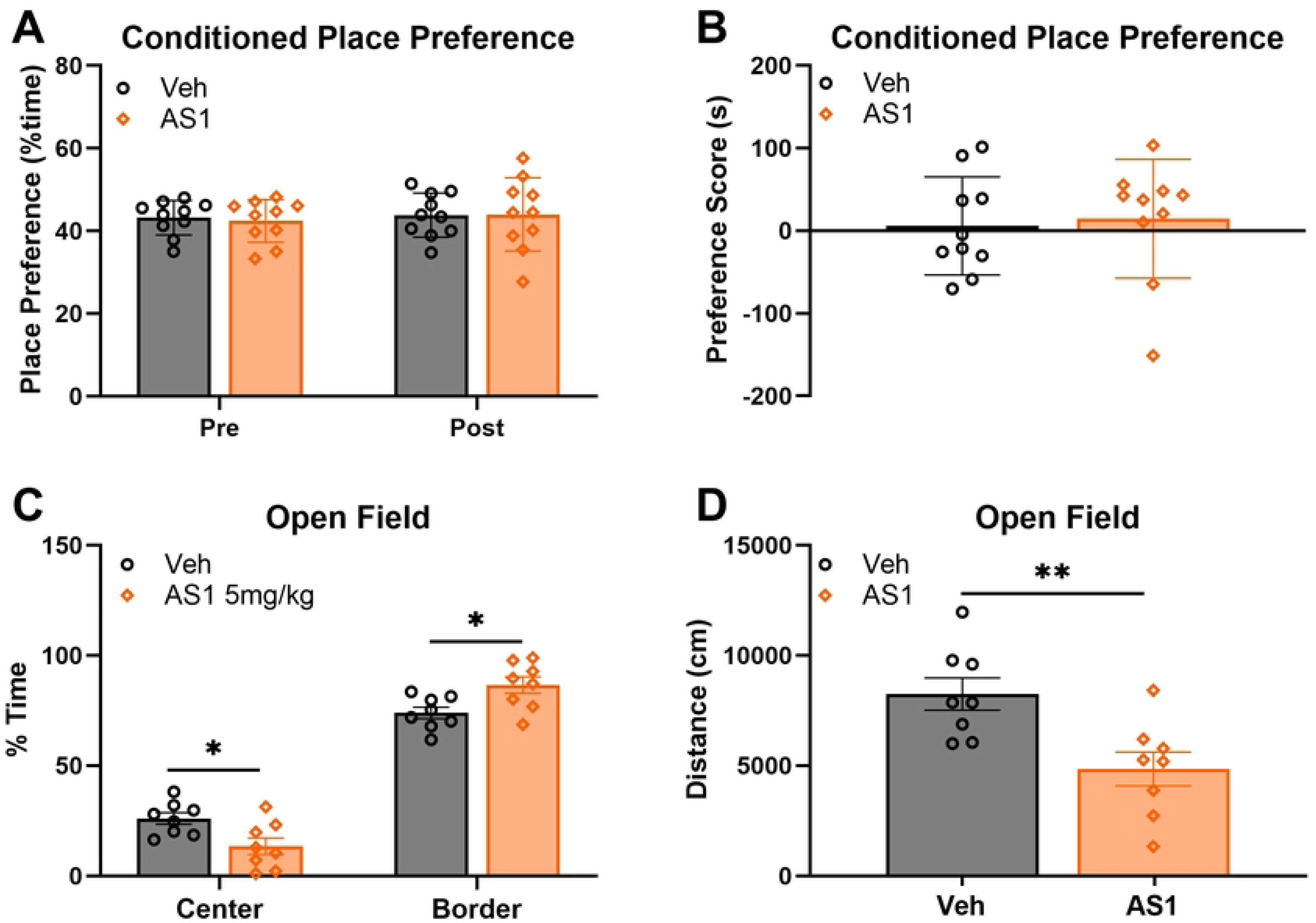
AS1 does not induce conditioned place preference but does reduce spontaneous locomotion in open field testing. (A,B) AS1 (5mg/kg) does alter place preference or preference score after conditioning (Veh n=10/5 females, AS1 n=10/5 females). B, AS1 (5mg/kg) inhibits locomotion and promotes time spent in the border during open field testing (Veh n=8/4 females, AS1 (5mg/kg) n=8/4 females). *p<0.05, **P<0.01. Veh=Vehicle, For=formalin. A-D, Unpaired student’s t-test.

### Open Field

To assess the effects of AS1 (5mg/kg) on spontaneous locomotor activity we employed open field testing. AS1 treated mice showed reduced occupancy in the center (AS1: 13.49 ± 3.74 vs Vehicle: 26.01 ±2.62, p=0.02) and increased occupancy at the border (AS1: 86.51 ± 3.74 vs Vehicle: 73.98 ±2.62 percent of time, p=0.02) when compared to vehicle treated mice (Figure 4C). AS1 treated mice also showed reduced total distance moved (AS1 4850.97 ±773.35 vs Vehicle: 8254.06 ±732.43 cm, p=0.006) (Fig 4D). Taken together, these results suggest that this dosage of AS1 reduced spontaneous locomotion (distance moved) which could be indicative of a sedative effect, and increased time at the border, which could be indicative of an anxiogenic effect (24).

### AS1 does not inhibit dorsal horn of the spinal cord activity evoked by formalin

Formalin evokes nociception by activating peripheral nociceptors which synapse with and excite second order nociceptive neurons in the dorsal horn of the spinal cord (21,25,26). To determine if AS1 acts to inhibit second phase formalin nocifensive behavior at the spinal cord level, we tested whether intrathecal administrated AS1 (5µg/µl) would affect formalin-induced licking behavior. Analysis revealed no effect of treatment (F(1, 15)=0.12, p=0.74). There were no differences between AS1 treated mice compared to the vehicle treated mice in both the first (AS1: 59.30 ±12.44 vs Vehicle: 98.00 ±16.35 sec, p=0.17) and second (AS1: 276.10 ±43.57 vs Vehicle: 213.00 ±36.00 sec, p=0.83) phases nor were there any individual timepoint differences (Fig 5A and 5B). This suggests that AS1 does not act at the level of the spinal cord to suppress formalin evoked nocifensive behavior.

**Fig 5.**
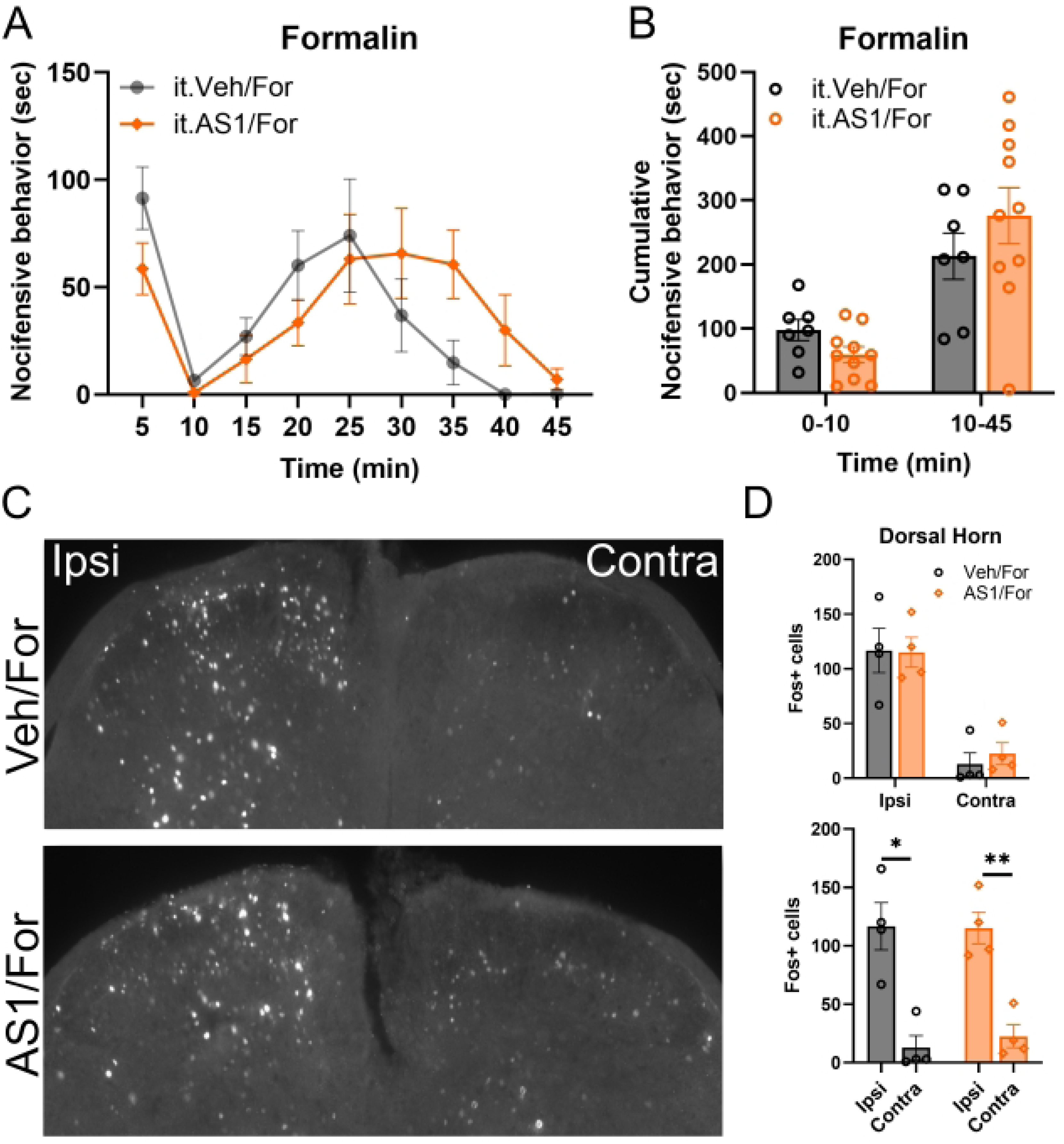
AS1 does not reduce formalin-evoked c-Fos immunoreactivity in the dorsal horn of the spinal cord. (A,B) Intrathecally administered AS1(5µg/µl) did not alter formalin evoked nocifensive behavior at any timepoint (A) or during phases one and two (B) (Veh i.t./For n=7/3 females, AS1 i.t./For n=10/3 females). (C) Formalin increased c-Fos immunoreactivity in the ipsilateral dorsal horn of the lumbar spinal cord but not in the contralateral dorsal horn. (D) Quantification of c-Fos positive neurons revealed no differences between vehicle or AS1-treated animals (top) but did reveal a significant increase in the number of c-Fos positive neurons in the ipsilateral dorsal horn compared to the contralateral dorsal horn for both conditions (bottom) (Veh n=4/2 females, AS1 n=4/2 females). *p<0.05, **P<0.01. Veh=Vehicle, For=formalin, ipsi=ipsilateral, contra=contralateral. A, RM-ANOVA with Bonferroni’s multiple comparison. B, Unpaired student t-teat. D, Unpaired student’s t-test (top), Paired student t-test (bottom).

To further explore this idea, we quantified c-Fos activity in the dorsal horn of the spinal cord following formalin treatment in the presence or absence of AS1 (5mg/kg, i.p.). We observed robust c-Fos staining in ipsilateral but not contralateral dorsal horn of both vehicle (p<0.05) and AS1 (p<0.01) treated animals with no difference between these treatment groups (AS1 ipsilateral: 115.25±13.68 neurons vs Vehicle ipsilateral: 116.75±20.24 neurons, p=0.95; AS1 contralateral: 22.5±9.77 neurons vs Vehicle contralateral: 12.75±10.43 neurons, p=0.17) (Fig 5C-D). Collectively, these data indicate that AS1 does not suppress formalin-evoked nocifensive behavior by suppressing primary nociceptor activation or activity in the dorsal horn of the spinal cord. This is consistent with findings observed in zebrafish where AS1 similarly did not affect primary nociceptor activity (10).

### AS1 increases c-Fos labelling in brain regions associated with pain processing

Neuronal activity profiling in larval zebrafish showed that AS1 in the presence of a noxious stimulus caused neuronal activity to increase in numerous regions including neuronal correlates of the nucleus accumbens, striatum, amygdala and the hypothalamus, which have all been implicated in pain processing (10). To determine if AS1 (5mg/kg, i.p.) might have similar properties in the mouse, we performed c-Fos neural activity expression profiling in formalin injected mice following either vehicle or AS1 administration and quantified the number of c-Fos positive cells in these regions. We found that compared to vehicle treated mice, the fold change (FC) of c-Fos positive cells were increased in the AS1 treated mice in the ACA (AS1: 5.45 ±1.14 vs Vehicle: 1.00 ±0.16, p=0.008), the shell of the NAc-shell (AS1: 2.83 ±0.37 vs Vehicle: 1.00 ±0.13, p=0.003), the PVH (AS1: 3.38 ±0.91 vs Vehicle: 1.00 ±0.33, p=0.04), the PVT (AS1: 3.40 ±0.55 vs Vehicle: 1.00 ±0.22, p=0.007), and the LS (AS1: 2.82 ±0.15 vs Vehicle: 1.00 ±0.27, p=0.001) (Fig 6). There were no fold change differences in the number of c-Fos positive cells between AS1 and vehicle treated mice in the NAc-Core (AS1: 2.38 ±0.16 vs Vehicle: 1.00 ±0.41, p=0.02), CeA (AS1: 1.79 ±0.29 vs 1.00 ±0.38, p=0.15), BLA (AS1: 1.85 ±0.13 vs Vehicle: 1.00 ±0.49, p=0.13), and MOp (AS1: 1.30 ±0.43 vs Vehicle: 1.00 ±0.42, p=0.63).

**Fig 6.**
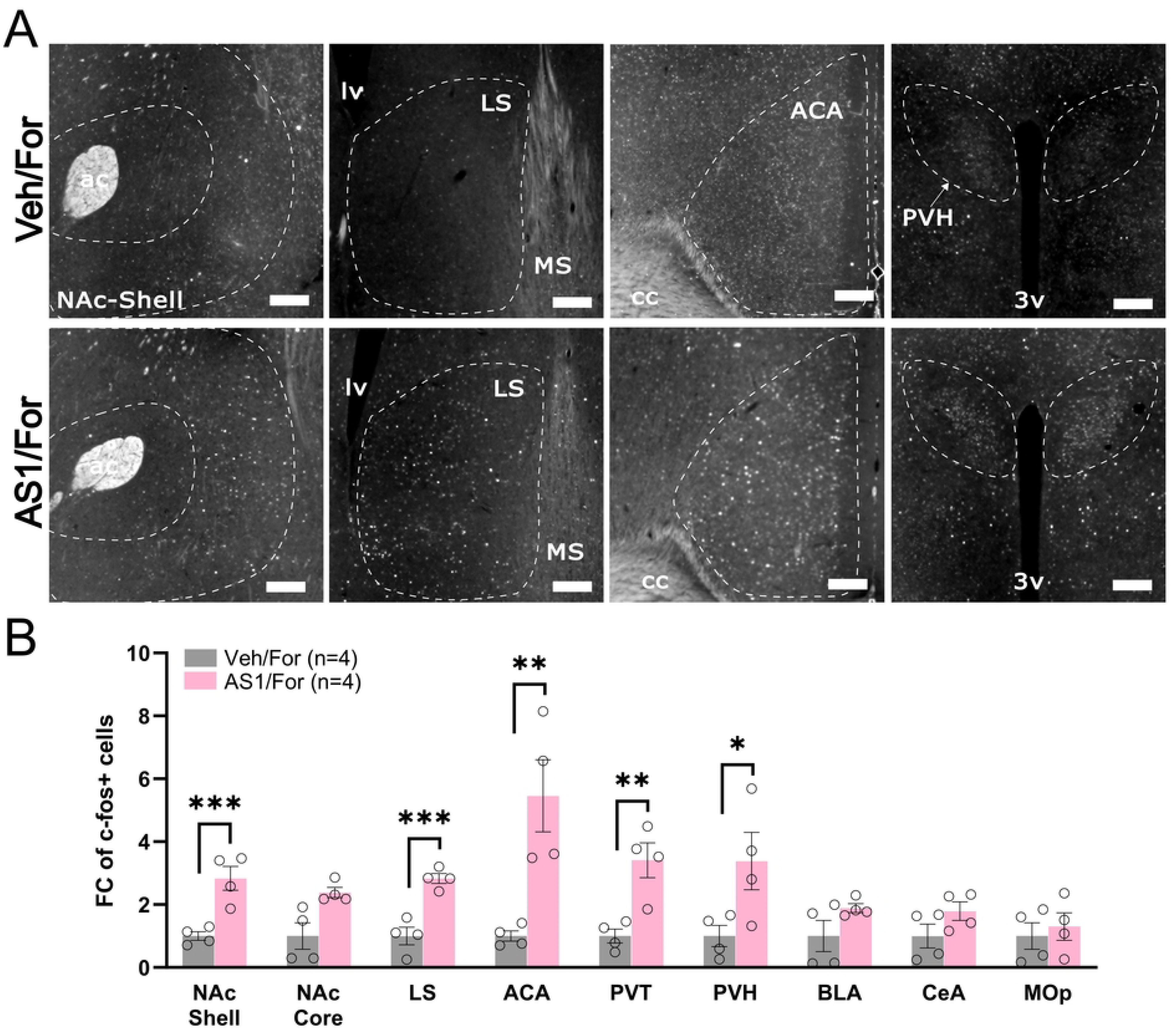
AS1 increases c-Fos immunoreactivity in brain regions of formalin treated animals associated with pain processing. (A) Representative images of c-Fos stained brain sections of vehicle/formalin (top) or AS1/formalin treated animals (veh/for n=4, AS1/for n=4). (B) Quantification of c-fos immunoreactivity. *p<0.05, **P<0.01, ***p<0.001. Veh=Vehicle, For=formalin, AP= anterior posterior; ACA= anterior cingulate area; cc= corpus callosum; ac= anterior commissure; NAc-Shell= nucleus accumbens shell; PVH=paraventricular hypothalamic nucleus; 3v= third ventricle; lv= lateral ventricle; LS= lateral septal nucleus; MS= medial septal nucleus. Scale bar: 100µm. B, Unpaired student’s t-test.

In a separate round of experiments, we investigated the neural activity induced by AS1 (5mg/kg) or vehicle with saline paw injection only to determine if any of the effects of AS1 were dependent on formalin treatment. We found that compared to vehicle treated mice, the fold change (FC) of c-fos positive cells were significantly increased in the AS1 treated mice in the ACA (AS1: 2.20 ±0.46 vs Vehicle: 1.00 ±0.08, p=0.04), CeA (AS1: 3.11 ±0.50 vs Vehicle: 1.00 ±0.19, p=0.007), and PVH (AS1: 1.77 ±0.22 vs Vehicle: 1.00 ±0.09, p=0.02) (**Fig**). There were no fold change differences in the number of c-fos positive cells between AS1 and vehicle treated mice in the NAc-Shell (AS1: 1.27 ±0.22 vs Vehicle: 1.00 ±0.10, p=0.30), NAc-Core (AS1: 1.94 ±0.42 vs Vehicle: 1.00 ±0.22, p=0.09), BLA (AS1: 1.63 ±0.28 vs Vehicle: 1.00 ± 0.14, p=0.09), LS (AS1: 1.15 ±0.14 vs Vehicle: 1.00 ±0.09, p=0.41), PVT (AS1: 1.27 ±0.10 vs Vehicle: 1.00 ±0.19, p=0.25), and MOp (AS1: 3.30 ±0.96 vs Vehicle: 1.00 ±0.50, p=0.07) (Fig 7). These data suggest that AS1 may alter brain activity in ways that are specific to formalin treatment. While c-Fos counts in the ACA and the PVH were elevated by AS1 treatment in both groups, only the NAc-shell, LS and PVT showed significantly higher c-Fos counts when AS1 was paired with formalin. Notably, fold change differences between AS1 and vehicle in the ACA and PVH were approximately two-fold greater in the formalin injected animals versus saline injected animals. In contrast, the CeA was only elevated when AS1 was paired with saline. Overall, these data suggest that the effects of AS1 on CNS activity are consistent with observation in larval zebrafish that some of these effects may be dependent on the presence of an aversive stimulus.

**Fig 7.**
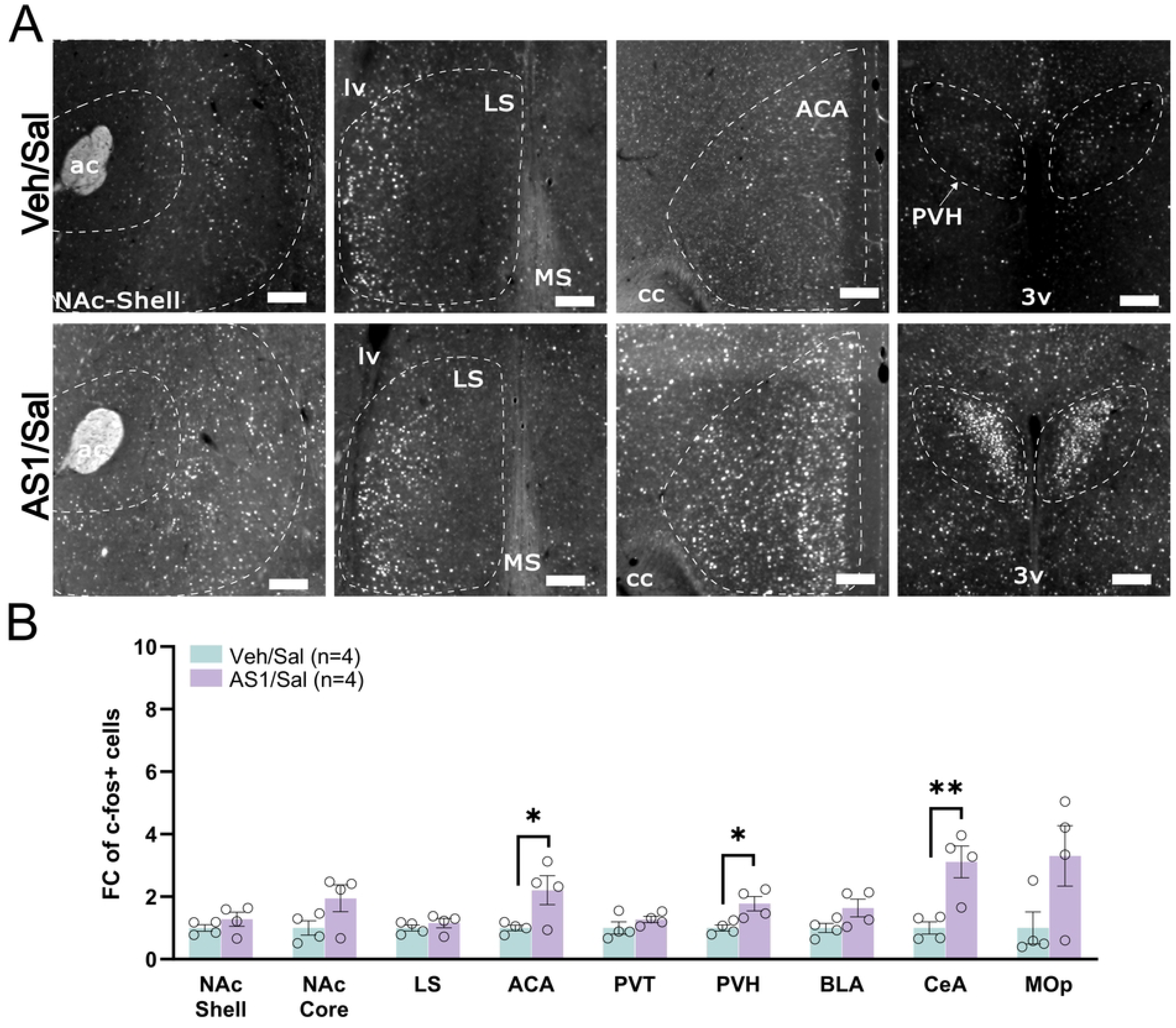
AS1 increases c-Fos immunoreactivity in brain regions, of saline treated animals, associated with pain processing. (A) Representative images of c-Fos stained brain sections of vehicle/saline (top) or AS1/saline treated animals (veh/sal n=4, AS1/sal n=4). (B) Quantification of c-fos immunoreactivity. *p<0.05, **P<0.01. Veh=Vehicle, sal=saline, AP= anterior posterior; ACA= anterior cingulate area; cc= corpus callosum; ac= anterior commissure; NAc-Shell= nucleus accumbens shell; PVH= paraventricular hypothalamic nucleus; 3v= third ventricle; lv= lateral ventricle; LS= lateral septal nucleus; MS= medial septal nucleus. Scale bar: 100µm

## Discussion

Here, we have described the effects of a small molecule AS1, that reverses the valence of noxious stimuli in larval zebrafish, in mouse models of nociception. We identified a dosage of AS1 that abolished second phase formalin evoked nocifensive behaviors, while having a limited effect on first phase behavior. AS1 had no effect on acute withdrawal responses to thermal and mechanical stimuli in the presence or absence of inflammation. Unlike in larval zebrafish, AS1 had no effect on thermal place preference. These data suggest that some of the analgesic properties of AS1 may be conserved in the mice, but not the ability of AS1 to reverse the valence of aversive stimuli.

The formalin assay is widely used to evaluate the analgesic potential of small of potential therapeutics. In this assay, formalin is injected into a hind paw. It is characterized by a first phase in which paw lifting, flicking and biting are observed. This phase lasts 5-10 minutes followed by a transient period where these behaviors subside. This period is followed by a second phase where these behaviors resume and subsides 45-60 minutes after formalin administration. The advantage of this assay is the spontaneous non-evoked nociceptive behavior can be observed in freely moving animals. Additionally, as behavior can be evaluated over a prolonged period, the time course of an agent’s effectiveness can be determined (20,22,27). The first phase is the result of primary nociceptor activity dependent on the noxious receptor TRPA1 whereas the second phase is thought to be dependent on both primary afferent activation and central sensitization (21,22,28). This second phase has long been thought to be driven by an inflammatory response, although a number of studies have called this assertion into question (29,30). Many compounds have been shown to differentially target the biphasic responses to formalin (31). Some drugs such as CBD, remifentanil, aspirin, and local anesthetics preferentially target the first phase, whereas others such as THC, morphine, bradykinin receptor antagonists, AMPA and NMDA antagonists target both phases. AS1 appears to fall into a group of compounds, with a variety of mechanisms of action, such as celecoxib, gabapentin, indomethacin, JNK inhibitors and adenosine which have been shown to preferentially target second phase responses (25,32–41).

Open field testing revealed that AS1 suppressed locomotion with a concurrent increase in the time spent in the border region. This could be indicative of a sedative effect which might contribute to the absence of second phase formalin response (24). This in itself would not necessarily rule out an analgesic effect of AS1, as sedative compounds such as diazepam have been found to have analgesic properties distinguishable from their sedative effects (42). Additionally, analgesic dosages of psychoactive compounds such as THC have also been found to impair locomotion and evoke anxiogenic behavior (13,43). Furthermore, the lack of an effect on first phase formalin and acute withdrawal responses to noxious stimuli suggests that AS1-treated animals have sufficient motor function to perform nociceptive behaviors which would argue for an analgesic property. Unlike diazepam which has been shown to produce conditioned place preference and thought to be rewarding, the AS1 dosing regimen employed in this study did not produce conditioned place preference or aversion (44). This suggests that this dosage of AS1 is neither rewarding nor aversive in the time period that AS1 evokes analgesic-like effects. This does not rule out that AS1 maybe be either rewarding or aversive if different dosing protocol was employed. Numerous compounds including cocaine and THC have been demonstrated to produce different outcomes in the CPP assay when this variable is altered (45).

The lack of effect of intrathecally-administered AS1 on formalin-evoked nocifensive behavior as well as the inability of AS1 to alter dorsal horn neuronal activity following formalin administration indicates that AS1 does not act on peripheral nociceptors or dorsal horn spinal neurons to mediate its effects. These findings align with previous findings in zebrafish where AS1 did not affect peripheral nociceptor activity. In larval zebrafish, AS1 in the presence of noxious stimuli led to increased neuronal activity in the neural correlates of the nucleus accumbens, striatum, hypothalamus, septal nuclei and the amygdala (10). Here, we observed similar results in the mouse, with an increase in c-Fos reactivity in the shell of the nucleus accumbens, the lateral septum, paraventricular thalamus, paraventricular hypothalamus and anterior cingulate cortex in AS1/formalin treated animals vs vehicle/formalin treated animals alone. We also observed a significant increase in neuronal activity in the anterior cingulate cortex and paraventricular hypothalamus of AS1/saline vs vehicle/saline animals however the foldchange difference was roughly two-fold less. As all these areas have been implicated in pain processing, it is conceivable that one or more of these areas could contribute to the observed effects of AS1 on second phase formalin responses (46–51).

In conclusion, AS1 appears to preferentially inhibit second phase formalin induced nocifensive behavior, but its hypolocomotor effects likely limit this compound’s utility as an analgesic. AS1 however may be a useful tool to interrogate the central mechanisms that facilitate second phase behavior. Future studies will be needed to identify the molecular and neuronal targets of this compound.

## Acknowledgements

We would like to thank Kyra Shelton for her assistance in performing experiments and Bryce Lecamp for thoughtful discussions.

